# Ownership, Coverage, Utilisation and Maintenance of Long-lasting insecticidal nets in Three Health Districts in Cameroon: A Cross-Sectional Study

**DOI:** 10.1101/465005

**Authors:** Frederick Nchang Cho, Paulette Ngum Fru, Blessing Menyi Cho, Solange Fri Munguh, Patrick Kofon Jokwi, Yayah Emerencia Ngah, Celestina Neh Fru, Andrew N Tassang, Albert Same Ekobo

## Abstract

The Bamenda, Santa and Tiko Health Districts are in the highest malaria transmission strata of Cameroon. The purpose of this study was to explore the indicators of ownership and utilisation as well as maintenance of long-lasting insecticidal nets (LLINs) in three health districts in Cameroon. A cross-sectional household survey involving 1,251 households was conducted in the Bamenda, Santa and Tiko Health Districts in Cameroon. A structured questionnaire was used to collect data on LLINs ownership, utilisation, and maintenance as well as demographic characteristics. The average number of LLINs per household was higher in the Bamenda Health District (BHD) compared to the Tiko Health District (THD) (2.5±1.4 vs 2.4±1.6) as well as the household ownership of at least one LLIN (93.3% vs. 88.9%). The proportion of the *de facto* population with universal utilisation was higher in BHD compared to THD (13.1% vs 0.2%). In multinomial regression analysis, households in the Santa Health District (SHD) (OR = 0.4, 95% = C.I; 0.2 – 0.8, *p* = 6.10×10^−3^), were less likely to own at least one LLIN, while those in the BHD (OR = 1.3, 95% = C.I; 0.8 – 2.1, *p* = 0.33) were more likely to maintain LLINs compared to those in THD. Ownership of LLINs was low in SHD and THD in comparison to the goal of one LLIN for every two household members. Overall, LLINs coverage and accessibility was still low after the free Mass Distribution Campaigns, making it difficult for all household members to effectively use LLINs.

## Introduction

Malaria is a preventable and curable disease transmitted by female Anopheles mosquitoes [1, 2] and it is a serious global public health problem with an estimated 216 million cases in 91 countries in 2016 [1, 3] and 2,000 cases in the United States each year [4]. Africa is the most a□ected region, with 90% of all estimated malaria cases and 91% of deaths in 2016 and 15 African countries contributing 80% of all cases [3, 5].

Cameroon, bordered by the Gulf of Guinea and Nigeria to the west; Chad and the Central African Republic to the east; and Equatorial Guinea, Gabon, and the Democratic Republic of Congo to the south, through the Ministry of Health (MOH), completed her third national universal long-lasting insecticidal net (LLIN) campaign in 2019. With support from the Global Fund, the MOH has made provision of free LLINs to pregnant women at antenatal care (ANC) clinics since 2008 [6, 7]. In 2011, the Cameroon MOH undertook a nationwide free LLINs distribution campaign from health facilities to all households, with the objective to provide a LLIN with a lifespan of five years, to all household beds or a LLIN for every two individuals per household, to a maximum of three LLINs per household [8, 9]. Malaria is endemic in Cameroon, with an estimated mortality rate of 11.6%, surpassing that of the African region of 10.4% as well as neighbouring countries [6]. It is the first major cause of morbidity and mortality among the most vulnerable groups [7, 10, 11]: children under five years and pregnant women, accounting, respectively, for 18% and 5% of the total population estimated at 19 million [12].

The main contemporary malaria control interventions are insecticide-treated bed nets and indoor residual spraying [2, 5, 7, 13, 14]. Alliance for Malaria Prevention has been instrumental in keeping long-lasting insecticide net (LLIN) campaigns on track: between 2014 and 2016 about 582 million LLINs were delivered globally and in 2017 there was the successful delivery of over 68 million nets to targeted recipients in Sub-Saharan Africa (SSA) and beyond [3]. Over 80% of all households have at least one mosquito net, up from 57% in 2011, still only about 60% of these households have enough nets to cover everyone at night [15]. The proportion of people in SSA sleeping under LLIN rose from less than 2% to over 50% between 2000-2015, preventing an estimated 450 million malaria cases [16].

Most studies in Cameroon and elsewhere have focused on various aspects of net ownership and utilisation. Effective LLIN use in the prevention of malaria in Mezam Division of the North West Region as well as the Tiko Health District in the South West Region [17, 18], *Plasmodium falciparum* infection in Rural and Semi-Urban Communities in the South West Region [8], predictive factors of ownership and utilisation in the Bamenda Health District (BHD) [19] and socio-demographic factors influencing the ownership and utilisation among malaria vulnerable groups in the Buea Health District and the Bamendankwe Health Area [20, 21]. However, there is paucity information on the indicators of LLIN ownership, utilisation, as well as maintenance. This study examines the indicators of net ownership and utilisation as well as maintenance, in three health districts in Cameroon.

## Materials and Methods

### Study design and setting

This cross-sectional survey conducted in the THD from June to July 2017 and in the BHD and SHD from March to May 2018 utilised a stratified multistage cluster sampling design. The study sampling frame included all health areas (HAs) in the study area, except those that were inaccessible for security reasons. Each Health Area (HA): urban, rural, or semi-urban localities were subdivided into quarters (primary sampling units or clusters in our study). On average, each HA had about five quarters. Sampled HAs in the THD had about 2,089 (35.58%) of the sampled population.

#### First stage

we randomly selected four HAs in the THD by probability proportionate to size (PPS) and conveniently sampled one each from the BHD and SHD.

#### Second stage

within each selected HA, we randomly selected at least three quarters and at most eight quarters by PPS (**Figure 1**).

**Figure 1:**
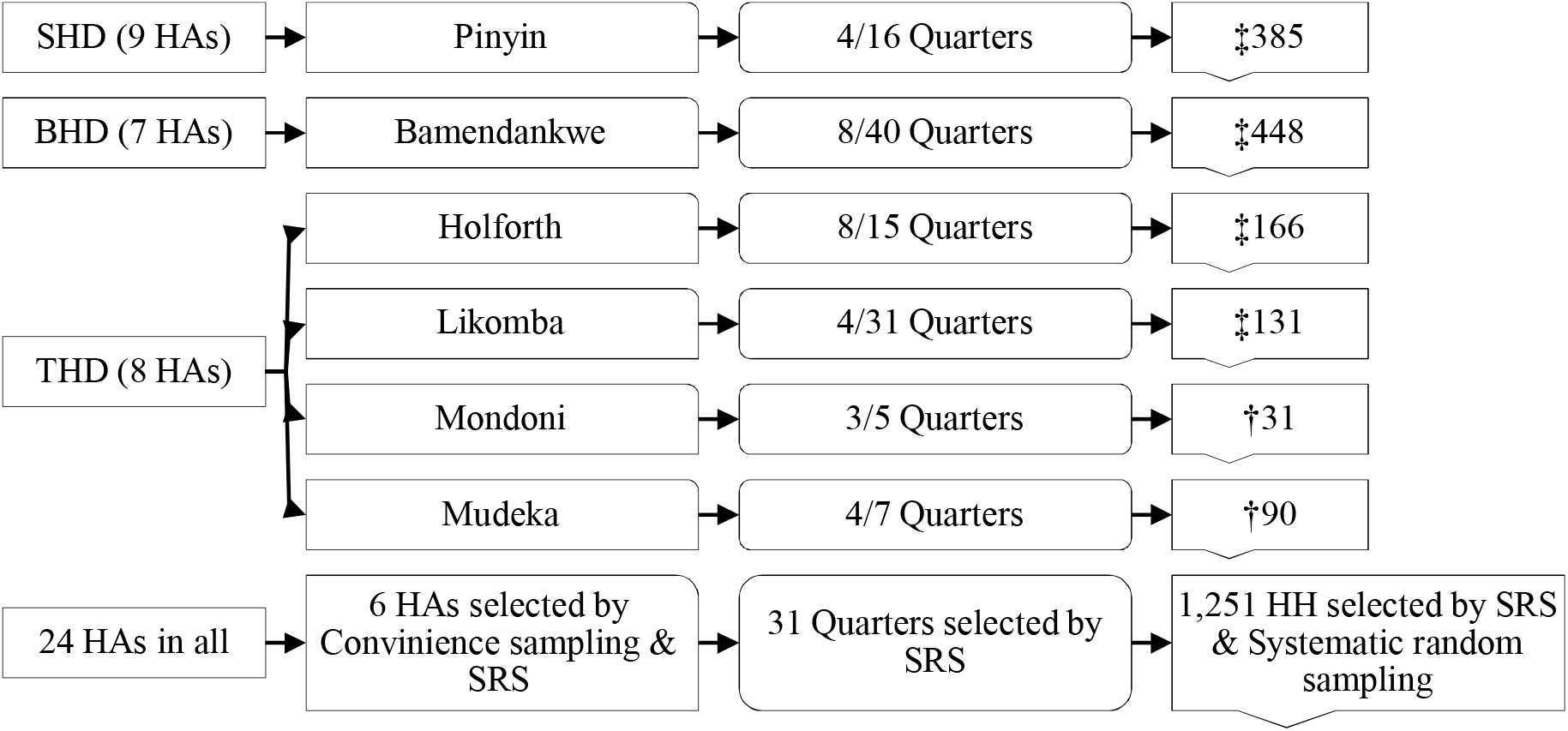
Multistage sampling (HA; Health Area, SRS; Simple Random Sampling) ‡Households (HH) sampled by systematic random sampling, †HHs sampled by SRS

#### Third stage

within each selected quarter, the survey team mapped and enumerated all households and selected households in each cluster by systematic random sampling (that is, a random start and interval to cover the entire quarter). The estimated number of households in each quarter was obtained from the quarter head to determine the sampling interval to select the households.

The study area consisted of the BHD with an estimated 350,000 residents and the Santa Health District (SHD), 35 Km from the BHD with 73,406 residents in the North West Region and the Tiko Health District (THD), 351 Km from the BHD with an estimated 134,649 residents in the South West Region of Cameroon [22]. Generally, malaria in Cameroon is caused mainly by *Plasmodium falciparum*, with *Anopheles gambiae* as the major vector [7, 12, 23]. The BHD (semi-urban community) and the SHD (rural community) are in the high western plateau altitude, where malaria transmission is permanent, occurring all year long, sometimes lessened by altitude although never totally absent [12, 24]. It is one of the most densely populated regions of Cameroon [7, 12]. The THD (urban and rural communities) is in the coastal strata, zone of dense hydrophile forest and mangrove swamp with the highest transmission of malaria in the country [7, 12]. Like the Buea health district, the THD has a constant variation in the trends of malaria prevalence all round the year [25, 26].

### Sample size determination

A minimum sample size of 384 for each health district, which was calculated with the CDC-Epi-Info version 7.2.2.6 (Centre for Disease Control, Georgia USA) StatCalc with the following characteristics, an average population of 307,620 in 2009 with an annual increase rate of 2% (6152.4) to 369,144 in 2018 [27], estimated proportion of households owning LLINs of 50%, accepted error margin of 5%, design effect of 1.0 and three clusters.

### Recruitment procedures and measures

Interviewers explained the purpose of the study and obtained signed/verbal informed consent from the head of the household or spouse. In cases where the household head was absent, any elderly person who has lived in the house for at least the last 12 months replaced her/him. The questionnaire for this study is as described in another study [18].

### Outcome variables

#### 1. LLINs ownership indicators

*Ownership of LLINs, Coverage*, and *Access to LLINs within the household* were defined as described in other studies [10, 28].

#### 2. LLIN utilisation indicators

*Household universal utilisation of LLINs* was defined as described in earlier studies [10, 28, 29]. *LLINs utilisation by the vulnerable population in the household* was defined as the proportion of children under five (or pregnant women) that slept under a LLIN the previous night [10]. *Regularly sleeping under bed nets*: household heads who reported habitually using nets daily [30]. *Slept under LLINs the previous night (SULPN)*: proportion of household heads who slept under a LLIN the previous night, as described in earlier studies [18, 21].

#### 3. Maintenance of LLINs

proportion of households in which the household heads admitted respecting the recommended washing frequency of LLINs.

#### 4. Independent variables (IV)

considered for association with LLINs ownership, utilisation, and maintenance were age, gender, marital status, education, occupation, health district, house type, and household composition.

### Statistical analysis

We entered data into MS-Excel (Microsoft Inc. 2016) and analysed with IBM-SPSS Statistics 25.0 for windows (IBM-SPSS Corp., Chicago USA). The Chi square (χ^*2*^) test was used to compare household characteristics with LLINs, ownership/utilisation indicators and multivariate logistic regression to identify significant correlates of the main outcomes. Statistical significance was set at *p* ≤ 0.05.

## Results

### Characteristics of the study participants

A total of 1,251 household heads was sampled with 5,870 *de facto* residents across six health areas in three health districts. Of the total household residents counted, 1,267 (21.6%) were children 0 – 5 years old and 93 (1.6%) were expectant mothers. The general characteristics of the study participants are shown in **Table 1**. The overall mean (±SD) family size was 4.7±2.1 members: 4.6±2.1 in BHD and 5.0±2.5 in THD. Majority of the houses, 1,060 (84.7%) were made of mud/cement, while households with 1 – 3 bedrooms; the mean (±SD) number of bedrooms 2.0±1.1 were about 1,141 (91.2%).

**Table 1:**
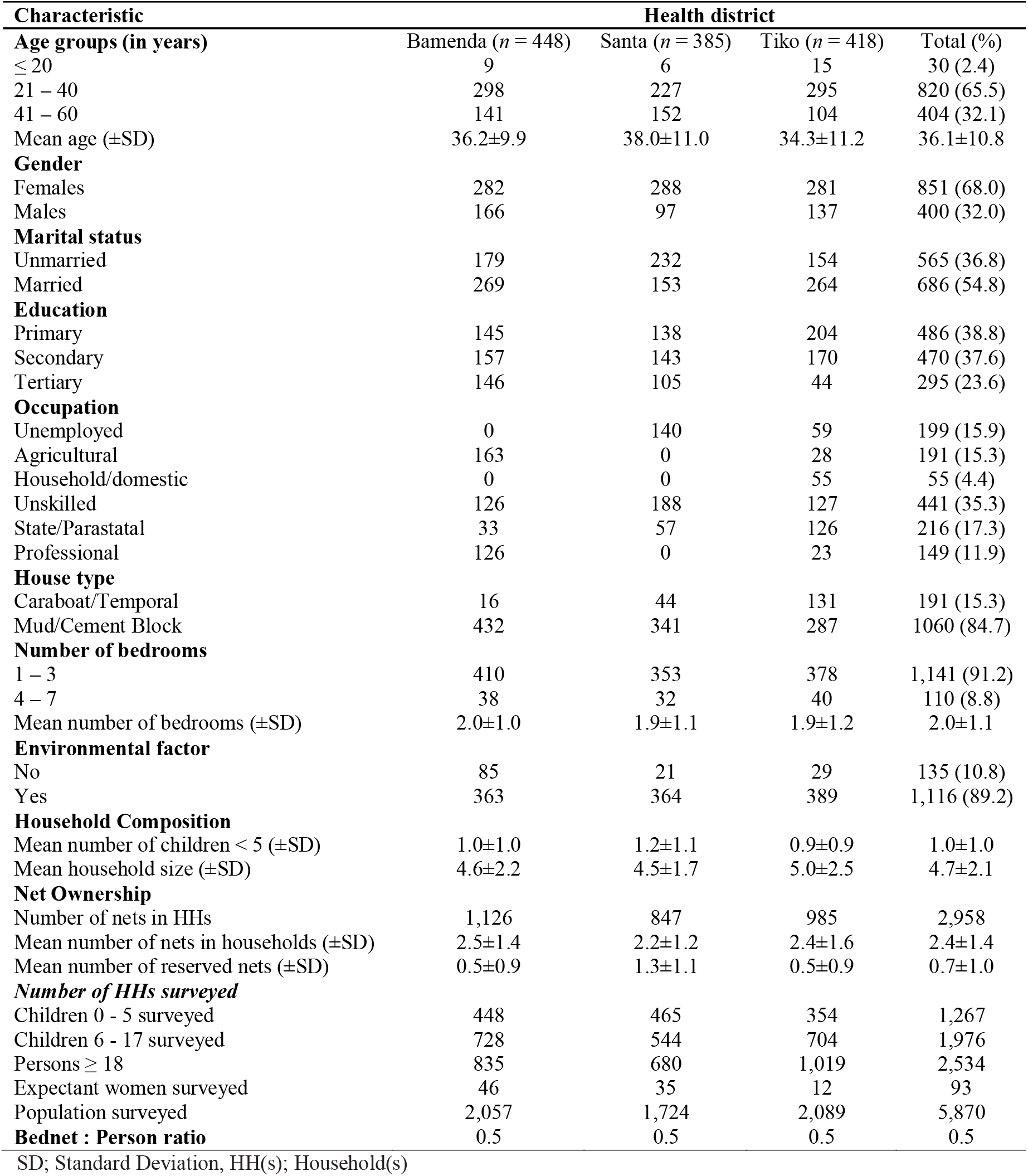
General characteristics of the study population.

### LLINs ownership, coverage and accessibility

One thousand one hundred and fifty-seven (92.5%) of the 1,251 households sampled had at least one LLIN, while 836 (66.8%) had at least a bednet for every two persons in the household (**Table 2**). The overall LLIN-to-person ratio was 0.50, that is, one net for every two persons (**Table 1**), constituting a coverage of 3,913 (66.7%) of the *de facto* population. The mean number of LLINs/LLINs density in the households was 2.4±1.4.

**Table 2:** LLINs ownership and utilisation indicators in association with health districts.

|  | Households |  |  |  |  |  | De facto population in households |  |  |  |
| --- | --- | --- | --- | --- | --- | --- | --- | --- | --- | --- |
| LLINs indicator | *n* (%) | BHD | SHD | THD | $\chi^2$ | *p*-value | *n* (%) | BHD | SHD | THD |
| **Ownership** |  |  |  |  |  |  |  |  |  |  |
| At least One | 1,157 (92.5) | 418 | 367 | 372 | 12.23 | **2.22x10<sup>-3</sup>** | 5,577 (95.0) | 2,000 | 1,680 | 1,897 |
| Coverage | 836 (66.8) | 387 | 214 | 235 | 120.46 | **6.97x10<sup>-27</sup>** | 3,913 (66.7) | 1,893 | 937 | 1,083 |
| Accessibility | 865 (69.1) | 374 | 214 | 277 | 77.97 | **6.97x10<sup>-27</sup>** | 4,058 (69.1) | 1,825 | 937 | 1,296 |
| **Utilisation** |  |  |  |  |  |  |  |  |  |  |
| Children 0 – 5 years | 558 (44.6) | 251 | 138 | 169 | 74.69 | **5.68x10<sup>-13</sup>** | 859 (14.6) | 427 | 188 | 244 |
| Entire household | 879 (70.3) | 380 | 193 | 306 | 268.21 | **3.22x10<sup>-44</sup>** | 942 (16.0) | 767 | 10 | 165 |

**Boldface numbers** indicate significant *p* values, BHD; Bamenda Health District, SHD; Santa Health District, THD; Tiko Health District, $\chi^2$ ; Chi square

Overall household accessibility to bednets was 865 (69.1%). Household accessibility (**Table 3**) to bednets was associated with gender, marital status, occupation of the household head, and health districts, where household residents in housed headed by females, unmarried, those with domestic chores and those in the BHD and SHD significantly (*p* < 0.05) had more access to LLINs than the other groups. In univariate analysis, these characteristics also showed significant associations with accessibility. Four thousand and fifty-eight (69.1%) of the *de facto* population, from 865 (69.1%) of the 1,251 households sampled, had access to LLINs.

**Table 3:**
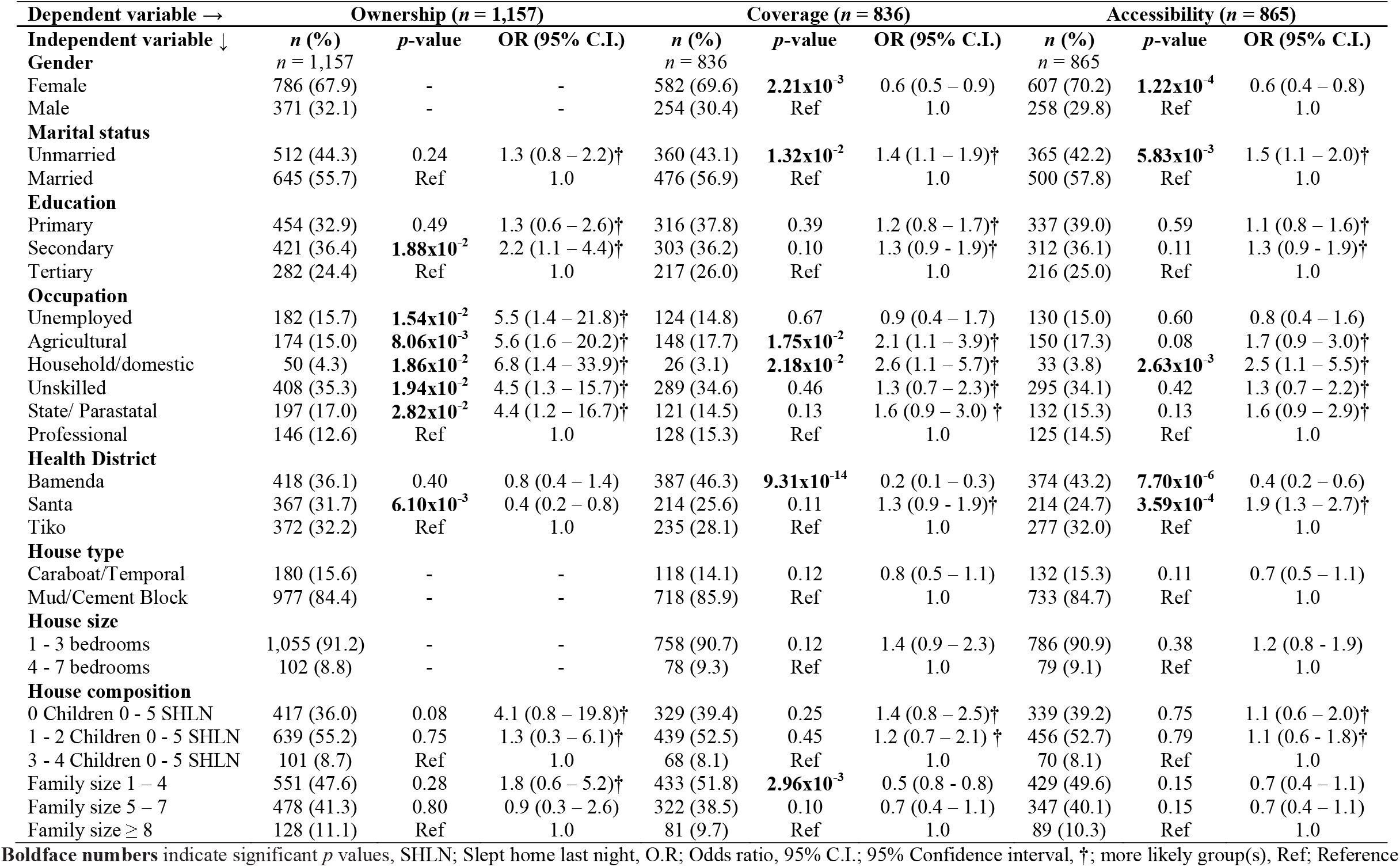
Multinomial logistic regression of socio-demographic characteristics in association with LLINs ownership indicators.

| Dependent variable → | Ownership (*n* = 1,157) |  |  | Coverage (*n* = 836) |  |  | Accessibility (*n* = 865) |  |  |
| --- | --- | --- | --- | --- | --- | --- | --- | --- | --- |
| Independent variable ↓ | *n* (%) | *p*-value | OR (95% C.I.) | *n* (%) | *p*-value | OR (95% C.I.) | *n* (%) | *p*-value | OR (95% C.I.) |
| **Gender** | *n* = 1,157 |  |  | *n* = 836 |  |  | *n* = 865 |  |  |
| Female | 786 (67.9) | - | - | 582 (69.6) | **2.21x10<sup>-3</sup>** | 0.6 (0.5 – 0.9) | 607 (70.2) | **1.22x10<sup>-4</sup>** | 0.6 (0.4 – 0.8) |
| Male | 371 (32.1) | - | - | 254 (30.4) | Ref | 1.0 | 258 (29.8) | Ref | 1.0 |
| **Marital status** |  |  |  |  |  |  |  |  |  |
| Unmarried | 512 (44.3) | 0.24 | 1.3 (0.8 – 2.2)† | 360 (43.1) | **1.32x10<sup>-2</sup>** | 1.4 (1.1 – 1.9)† | 365 (42.2) | **5.83x10<sup>-3</sup>** | 1.5 (1.1 – 2.0)† |
| Married | 645 (55.7) | Ref | 1.0 | 476 (56.9) | Ref | 1.0 | 500 (57.8) | Ref | 1.0 |
| **Education** |  |  |  |  |  |  |  |  |  |
| Primary | 454 (32.9) | 0.49 | 1.3 (0.6 – 2.6)† | 316 (37.8) | 0.39 | 1.2 (0.8 – 1.7)† | 337 (39.0) | 0.59 | 1.1 (0.8 – 1.6)† |
| Secondary | 421 (36.4) | **1.88x10<sup>-2</sup>** | 2.2 (1.1 – 4.4)† | 303 (36.2) | 0.10 | 1.3 (0.9 – 1.9)† | 312 (36.1) | 0.11 | 1.3 (0.9 – 1.9)† |
| Tertiary | 282 (24.4) | Ref | 1.0 | 217 (26.0) | Ref | 1.0 | 216 (25.0) | Ref | 1.0 |
| **Occupation** |  |  |  |  |  |  |  |  |  |
| Unemployed | 182 (15.7) | **1.54x10<sup>-2</sup>** | 5.5 (1.4 – 21.8)† | 124 (14.8) | 0.67 | 0.9 (0.4 – 1.7) | 130 (15.0) | 0.60 | 0.8 (0.4 – 1.6) |
| Agricultural | 174 (15.0) | **8.06x10<sup>-3</sup>** | 5.6 (1.6 – 20.2)† | 148 (17.7) | **1.75x10<sup>-2</sup>** | 2.1 (1.1 – 3.9)† | 150 (17.3) | 0.08 | 1.7 (0.9 – 3.0)† |
| Household/domestic | 50 (4.3) | **1.86x10<sup>-2</sup>** | 6.8 (1.4 – 33.9)† | 26 (3.1) | **2.18x10<sup>-2</sup>** | 2.6 (1.1 – 5.7)† | 33 (3.8) | **2.63x10<sup>-3</sup>** | 2.5 (1.1 – 5.5)† |
| Unskilled | 408 (35.3) | **1.94x10<sup>-2</sup>** | 4.5 (1.3 – 15.7)† | 289 (34.6) | 0.46 | 1.3 (0.7 – 2.3)† | 295 (34.1) | 0.42 | 1.3 (0.7 – 2.2)† |
| State/ Parastatal | 197 (17.0) | **2.82x10<sup>-2</sup>** | 4.4 (1.2 – 16.7)† | 121 (14.5) | 0.13 | 1.6 (0.9 – 3.0)† | 132 (15.3) | 0.13 | 1.6 (0.9 – 2.9)† |
| Professional | 146 (12.6) | Ref | 1.0 | 128 (15.3) | Ref | 1.0 | 125 (14.5) | Ref | 1.0 |
| **Health District** |  |  |  |  |  |  |  |  |  |
| Bamenda | 418 (36.1) | 0.40 | 0.8 (0.4 – 1.4) | 387 (46.3) | **9.31x10<sup>-14</sup>** | 0.2 (0.1 – 0.3) | 374 (43.2) | **7.70x10<sup>-6</sup>** | 0.4 (0.2 – 0.6) |
| Santa | 367 (31.7) | **6.10x10<sup>-3</sup>** | 0.4 (0.2 – 0.8) | 214 (25.6) | 0.11 | 1.3 (0.9 – 1.9)† | 214 (24.7) | **3.59x10<sup>-4</sup>** | 1.9 (1.3 – 2.7)† |
| Tiko | 372 (32.2) | Ref | 1.0 | 235 (28.1) | Ref | 1.0 | 277 (32.0) | Ref | 1.0 |
| **House type** |  |  |  |  |  |  |  |  |  |
| Caraboat/Temporal | 180 (15.6) | - | - | 118 (14.1) | 0.12 | 0.8 (0.5 – 1.1) | 132 (15.3) | 0.11 | 0.7 (0.5 – 1.1) |
| Mud/Cement Block | 977 (84.4) | - | - | 718 (85.9) | Ref | 1.0 | 733 (84.7) | Ref | 1.0 |
| **House size** |  |  |  |  |  |  |  |  |  |
| 1 - 3 bedrooms | 1,055 (91.2) | - | - | 758 (90.7) | 0.12 | 1.4 (0.9 – 2.3) | 786 (90.9) | 0.38 | 1.2 (0.8 – 1.9) |
| 4 - 7 bedrooms | 102 (8.8) | - | - | 78 (9.3) | Ref | 1.0 | 79 (9.1) | Ref | 1.0 |
| **House composition** |  |  |  |  |  |  |  |  |  |
| 0 Children 0 - 5 SHLN | 417 (36.0) | 0.08 | 4.1 (0.8 – 19.8)† | 329 (39.4) | 0.25 | 1.4 (0.8 – 2.5)† | 339 (39.2) | 0.75 | 1.1 (0.6 – 2.0)† |
| 1 - 2 Children 0 - 5 SHLN | 639 (55.2) | 0.75 | 1.3 (0.3 – 6.1)† | 439 (52.5) | 0.45 | 1.2 (0.7 – 2.1)† | 456 (52.7) | 0.79 | 1.1 (0.6 – 1.8)† |
| 3 - 4 Children 0 - 5 SHLN | 101 (8.7) | Ref | 1.0 | 68 (8.1) | Ref | 1.0 | 70 (8.1) | Ref | 1.0 |
| Family size 1 – 4 | 551 (47.6) | 0.28 | 1.8 (0.6 – 5.2)† | 433 (51.8) | **2.96x10<sup>-3</sup>** | 0.5 (0.8 – 0.8) | 429 (49.6) | 0.15 | 0.7 (0.4 – 1.1) |
| Family size 5 – 7 | 478 (41.3) | 0.80 | 0.9 (0.3 – 2.6) | 322 (38.5) | 0.10 | 0.7 (0.4 – 1.1) | 347 (40.1) | 0.15 | 0.7 (0.4 – 1.1) |
| Family size ≥ 8 | 128 (11.1) | Ref | 1.0 | 81 (9.7) | Ref | 1.0 | 89 (10.3) | Ref | 1.0 |

**Boldface numbers** indicate significant *p* values, SHLN; Slept home last night, O.R; Odds ratio, 95% C.I.; 95% Confidence interval, †; more likely group(s), Ref; Reference

In univariate and multivariate analyses, LLINs ownership, coverage and accessibility were associated (*p* < 0.05) with health districts (**Table 3**), where households in the SHD significantly (*p* = 6.1×10^−3^) owned fewer nets, while those in the BHD significantly (*p* = 9.31×10^−14^) had more coverage than the other districts. Ownership, coverage and accessibility were associated with marital status both in univariate and multivariate analysis. Coverage was also associated with the gender of the household head and household size (**Table 3**), where households headed by females (*p* = 2.21×10^−3^) and those of household size of 1 – 4 members (*p* = 2.9×10^−3^) significantly influenced coverage than the others. Secondary educational and occupational status significantly influenced household ownership of nets (*p* < 0.05). Household heads in the agricultural sector (OR = 2.1, 95% C.I = 1.1 – 3.9, *p* = 1.75×10^−2^) as well as those in domestic affairs (OR = 2.6, 95% C.I = 1.1 –5.7, *p* = 2.18×10^−2^), were more likely to have LLINs coverage when compared with their counterparts (**Table 3**).

### Use of LLINs

Of the 1,251 households sampled, 520 (41.6%) and 256 (20.5%) were those in which all children 0 – 5 years and those in which all who slept home last night used bednets, respectively representing 859 (14.6%) and 942 (16.0%) of the 5,870 *de facto* population that slept home last night (**Table 2**).

Marital status (OR = 2.2, 95% C.I = 1.5 – 3.2, *p* = 6.09×10^−5^), vs (OR = 1.4, 95% C.I = 1.0 – 2.1, *p* = 4.40×10^−2^), household size (OR = 1.3, 95% C.I = 0.6 – 2.7, *p* = 0.45) vs (OR = 1.2, 95% C.I = 0.7 – 2.2) and the installation of LLINs on all beds in the household were the predictors of the utilisation of LLINs by all children 0 – 5 years old and the entire household (**Table 4**).

**Table 4:**
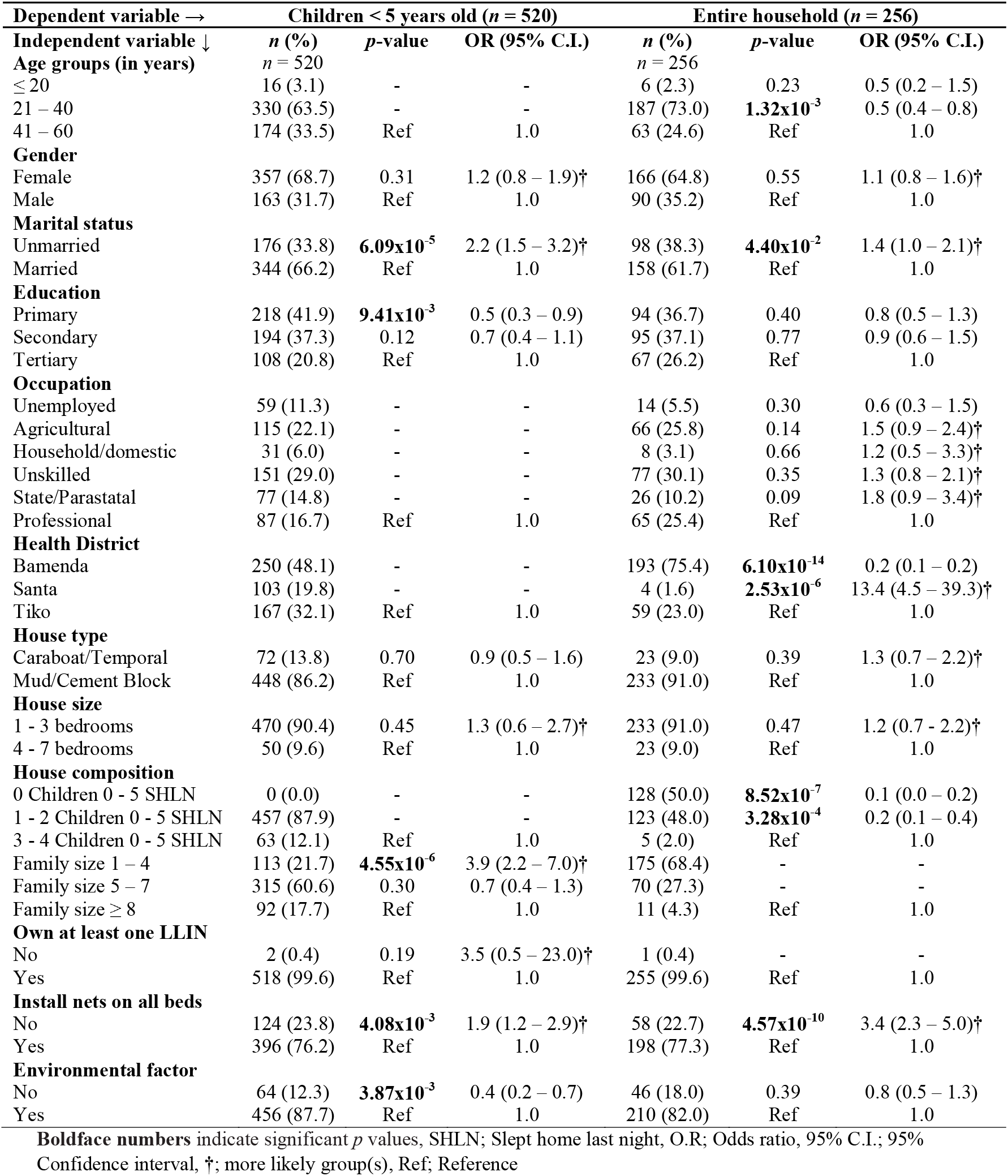
Multinomial logistic regression of socio-demographic characteristics in association with LLINs use by all children 0 – 5 and the entire household.

| Dependent variable → | Children < 5 years old (*n* = 520) |  |  | Entire household (*n* = 256) |  |  |
| --- | --- | --- | --- | --- | --- | --- |
| Independent variable ↓ | *n* (%) | *p*-value | OR (95% C.I.) | *n* (%) | *p*-value | OR (95% C.I.) |
| **Age groups (in years)** | *n* = 520 |  |  | *n* = 256 |  |  |
| ≤ 20 | 16 (3.1) | - | - | 6 (2.3) | 0.23 | 0.5 (0.2 – 1.5) |
| 21 – 40 | 330 (63.5) | - | - | 187 (73.0) | **1.32x10<sup>-3</sup>** | 0.5 (0.4 – 0.8) |
| 41 – 60 | 174 (33.5) | Ref | 1.0 | 63 (24.6) | Ref | 1.0 |
| **Gender** |  |  |  |  |  |  |
| Female | 357 (68.7) | 0.31 | 1.2 (0.8 – 1.9)† | 166 (64.8) | 0.55 | 1.1 (0.8 – 1.6)† |
| Male | 163 (31.7) | Ref | 1.0 | 90 (35.2) | Ref | 1.0 |
| **Marital status** |  |  |  |  |  |  |
| Unmarried | 176 (33.8) | **6.09x10<sup>-5</sup>** | 2.2 (1.5 – 3.2)† | 98 (38.3) | **4.40x10<sup>-2</sup>** | 1.4 (1.0 – 2.1)† |
| Married | 344 (66.2) | Ref | 1.0 | 158 (61.7) | Ref | 1.0 |
| **Education** |  |  |  |  |  |  |
| Primary | 218 (41.9) | **9.41x10<sup>-3</sup>** | 0.5 (0.3 – 0.9) | 94 (36.7) | 0.40 | 0.8 (0.5 – 1.3) |
| Secondary | 194 (37.3) | 0.12 | 0.7 (0.4 – 1.1) | 95 (37.1) | 0.77 | 0.9 (0.6 – 1.5) |
| Tertiary | 108 (20.8) | Ref | 1.0 | 67 (26.2) | Ref | 1.0 |
| **Occupation** |  |  |  |  |  |  |
| Unemployed | 59 (11.3) | - | - | 14 (5.5) | 0.30 | 0.6 (0.3 – 1.5) |
| Agricultural | 115 (22.1) | - | - | 66 (25.8) | 0.14 | 1.5 (0.9 – 2.4)† |
| Household/domestic | 31 (6.0) | - | - | 8 (3.1) | 0.66 | 1.2 (0.5 – 3.3)† |
| Unskilled | 151 (29.0) | - | - | 77 (30.1) | 0.35 | 1.3 (0.8 – 2.1)† |
| State/Parastatal | 77 (14.8) | - | - | 26 (10.2) | 0.09 | 1.8 (0.9 – 3.4)† |
| Professional | 87 (16.7) | Ref | 1.0 | 65 (25.4) | Ref | 1.0 |
| **Health District** |  |  |  |  |  |  |
| Bamenda | 250 (48.1) | - | - | 193 (75.4) | **6.10x10<sup>-14</sup>** | 0.2 (0.1 – 0.2) |
| Santa | 103 (19.8) | - | - | 4 (1.6) | **2.53x10<sup>-6</sup>** | 13.4 (4.5 – 39.3)† |
| Tiko | 167 (32.1) | Ref | 1.0 | 59 (23.0) | Ref | 1.0 |
| **House type** |  |  |  |  |  |  |
| Caraboat/Temporal | 72 (13.8) | 0.70 | 0.9 (0.5 – 1.6) | 23 (9.0) | 0.39 | 1.3 (0.7 – 2.2)† |
| Mud/Cement Block | 448 (86.2) | Ref | 1.0 | 233 (91.0) | Ref | 1.0 |
| **House size** |  |  |  |  |  |  |
| 1 - 3 bedrooms | 470 (90.4) | 0.45 | 1.3 (0.6 – 2.7)† | 233 (91.0) | 0.47 | 1.2 (0.7 - 2.2)† |
| 4 - 7 bedrooms | 50 (9.6) | Ref | 1.0 | 23 (9.0) | Ref | 1.0 |
| **House composition** |  |  |  |  |  |  |
| 0 Children 0 - 5 SHLN | 0 (0.0) | - | - | 128 (50.0) | **8.52x10<sup>-7</sup>** | 0.1 (0.0 – 0.2) |
| 1 - 2 Children 0 - 5 SHLN | 457 (87.9) | - | - | 123 (48.0) | **3.28x10<sup>-4</sup>** | 0.2 (0.1 – 0.4) |
| 3 - 4 Children 0 - 5 SHLN | 63 (12.1) | Ref | 1.0 | 5 (2.0) | Ref | 1.0 |
| Family size 1 – 4 | 113 (21.7) | **4.55x10<sup>-6</sup>** | 3.9 (2.2 – 7.0)† | 175 (68.4) | - | - |
| Family size 5 – 7 | 315 (60.6) | 0.30 | 0.7 (0.4 – 1.3) | 70 (27.3) | - | - |
| Family size ≥ 8 | 92 (17.7) | Ref | 1.0 | 11 (4.3) | Ref | 1.0 |
| **Own at least one LLIN** |  |  |  |  |  |  |
| No | 2 (0.4) | 0.19 | 3.5 (0.5 – 23.0)† | 1 (0.4) | - | - |
| Yes | 518 (99.6) | Ref | 1.0 | 255 (99.6) | Ref | 1.0 |
| **Install nets on all beds** |  |  |  |  |  |  |
| No | 124 (23.8) | **4.08x10<sup>-3</sup>** | 1.9 (1.2 – 2.9)† | 58 (22.7) | **4.57x10<sup>-10</sup>** | 3.4 (2.3 – 5.0)† |
| Yes | 396 (76.2) | Ref | 1.0 | 198 (77.3) | Ref | 1.0 |
| **Environmental factor** |  |  |  |  |  |  |
| No | 64 (12.3) | **3.87x10<sup>-3</sup>** | 0.4 (0.2 – 0.7) | 46 (18.0) | 0.39 | 0.8 (0.5 – 1.3) |
| Yes | 456 (87.7) | Ref | 1.0 | 210 (82.0) | Ref | 1.0 |

**Boldface numbers** indicate significant *p* values, SHLN; Slept home last night, O.R; Odds ratio, 95% C.I.; 95% Confidence interval, †; more likely group(s), Ref; Reference

Of the 1,251 household heads sampled, 484 (38.7%) regularly used bednets on all nights of the week, while 350 (28.0%) had their household heads using bednets last night (**Table 2**).

The other uses, “out of norms”, of LLINs are summarised in **Table 5**. Three hundred and fifty-nine [359 (28.7%); 95% C.I = 26.3 – 31.3] of the household heads sampled, admitted that LLINs were put into other diverse uses. These uses ranged from being used as goal post nets by children; 2.8% (95% C.I = 2.0 – 3.9), to yard fences; 22.7% (95% C.I = 20.5 – 25.1). Except for harvesting and drying of melon seeds (egussi), the other “out of norm” uses of LLINs were significantly (*p* < 0.05) associated to the health districts.

**Table 5:**
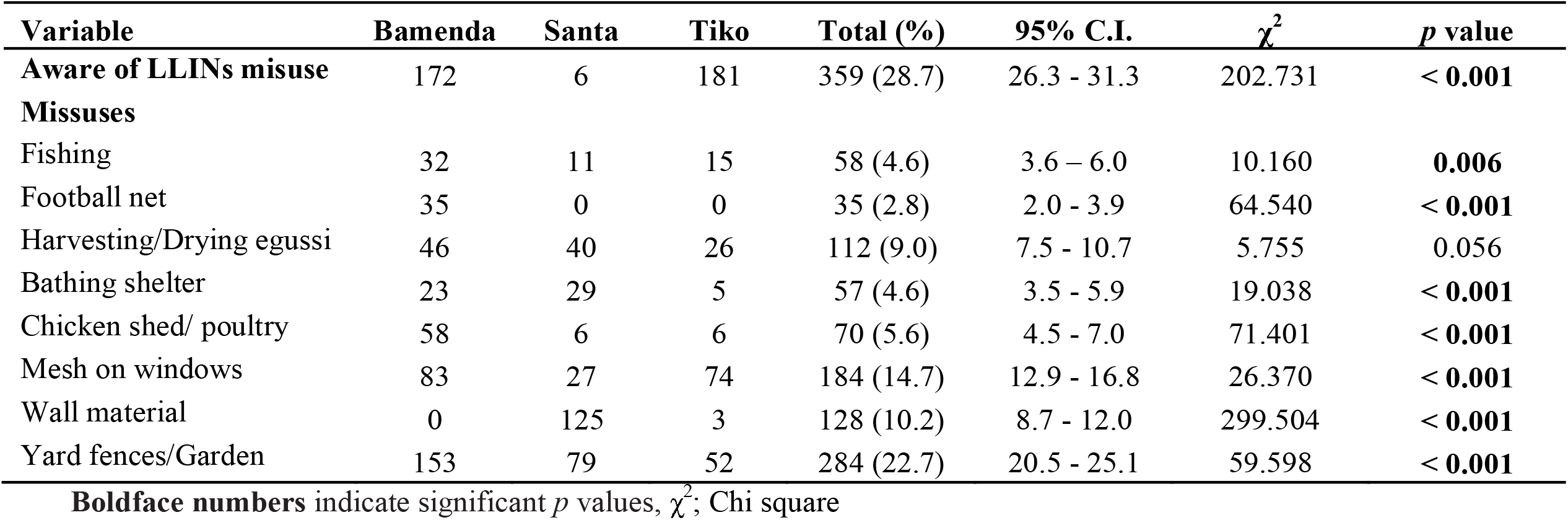
LLINs Misappropriation: by health districts.

### Care and maintenance of LLINs

Out of the 1,251 household heads sampled, 1,089 (87.1%) said LLINs can be washed, while 627 (50.1%) affirmed the recommended LLINs washing frequency of 2 – 4 times a year (**Table 6**).

**Table 6:** Logistic regression of socio-demographic characteristics in association with LLINs maintenance.

| Dependent variable → | Can wash LLINs (n = 1,089) |  |  | Recommended wash frequency (n = 627) |  |  |
| --- | --- | --- | --- | --- | --- | --- |
| Independent variable ↓ | n (%) | p-value | OR (95% C.I.) | n (%) | p-value | OR (95% C.I.) |
| **Age groups (in years)** | n = 1,089 |  |  | n = 627 |  |  |
| ≤ 20 | 24 (2.2) | 0.20 | 1.9 (0.7 – 5.0)† | 10 (1.6) | 0.25 | 1.6 (0.7 – 3.7)† |
| 21 – 40 | 709 (65.1) | 0.16 | 1.3 (0.9 - 1.9) † | 424 (67.6) | 0.25 | 0.9 (0.7 - 1.1) |
| 41 – 60 | 356 (32.7) | Ref | 1.0 | 193 (30.8) | Ref | 1.0 |
| **Gender** |  |  |  |  |  |  |
| Female | 775 (69.3) | **2.22x10<sup>-2</sup>** | 0.7 (0.5 – 0.9) | 431 (68.7) | 0.08 | 0.8 (0.6 - 1.0) |
| Male | 334 (30.7) | Ref | 1.0 | 196 (31.3) | Ref | 1.0 |
| **Marital status** |  |  |  |  |  |  |
| Unmarried | 485 (44.5) | 0.63 | 1.1 (0.8 - 1.6)† | 263 (41.9) | 0.16 | 1.2 (0.9 - 1.5)† |
| Married | 604 (55.5) | Ref | 1.0 | 364 (58.1) | Ref | 1.0 |
| **Education** |  |  |  |  |  |  |
| Primary | 422 (38.8) | 0.91 | 1.0 (0.6 – 1.5) | 233 (37.2) | 0.95 | 1.0 (0.7 - 1.4) |
| Secondary | 416 (38.2) | 0.20 | 0.7 (0.5 - 1.2) | 235 (37.5) | 0.77 | 1.0 (0.8 - 1.3) |
| Tertiary | 251 (23.0) | Ref | 1.0 | 159 (25.4) | Ref | 1.0 |
| **Occupation** |  |  |  |  |  |  |
| Unemployed | 167 (15.3) | 0.23 | 1.6 (0.7 – 3.3)† | 72 (11.5) | 0.19 | 1.4 (0.8 – 2.5)† |
| Agricultural | 163 (15.0) | 0.92 | 1.0 (0.5 - 1.8) | 133 (21.2) | 0.48 | 0.8 (0.5 – 1.4) |
| Household/domestic | 54 (5.0) | 0.07 | 0.1 (0.0 – 1.2) | 19 (3.0) | 0.06 | 2.0 (1.0 – 4.2)† |
| Unskilled | 389 (35.7) | 0.94 | 1.0 (0.5 – 1.8) | 202 (32.2) | 0.19 | 1.3 (0.9 – 2.1)† |
| State/Parastatal | 190 (17.4) | 0.80 | 0.9 (0.5 – 1.8) | 104 (16.6) | 0.79 | 1.1 (0.7 - 1.8)† |
| Professional | 126 (11.6) | Ref | 1.0 | 97 (15.5) | Ref | 1.0 |
| **Health District** |  |  |  |  |  |  |
| Bamenda | 379 (34.8) | 0.33 | 1.3 (0.8 – 2.1)† | 306 (48.8) | **3.14x10<sup>-7</sup>** | 0.4 (0.3 – 0.6) |
| Santa | 341 (31.3) | 0.35 | 0.8 (0.5 - 1.3) | 146 (23.3) | 0.61 | 1.1 (0.8 - 1.5)† |
| Tiko | 369 (33.9) | Ref | 1.0 | 175 (27.9) | Ref | 1.0 |

**Boldface numbers** indicate significant *p* values, O.R; Odds ratio, 95% C.I.; 95% Confidence interval, †; more likely group(s), Ref; Reference

The question of washing bednets or not, was associated to the gender of the household head, where households with females heads significantly (*p* = 2.22×10^−2^) washed them compared to those headed by males.

The WHO recommended washing frequency was associated with the health district, where households with heads in the BHD (*p* = 3.14×10^−7^) abided more significantly to the recommended washing frequency than those in the other health districts.

## DISCUSSION

This study examined the indicators of LLIN ownership, utilisation, and maintenance in the Bamenda, Santa and Tiko Health Districts. Overall, 92.5% and 20.5% of the households interviewed owned at least one LLIN per household and utilisation by the entire household last night, respectively.

### Indicators of household LLINs ownership

Currently, targets in national strategic plans for all three LLINs coverage indicators are usually set for all people at risk of malaria [1, 5], to ≥ 80%. Household ownership of at least one LLIN per household in this study is higher than rates reported elsewhere in and out of Cameroon [8, 17, 19, 20, 30-44]. It was however, lower than the proportions reported within the THD, Uganda, Ethiopia, and Myanmar [18, 45-47] and in line with the 93.3 – 98% reported in the Bamendankwe Health area, Madagascar and India [21, 33, 48]. The high proportion of owning at least a LLIN per household in this study could be attributed to the free LLINs MDCs [8, 12, 49].

The universal household coverage of 66.8%, although within the WHO range of 39 – 75 % [50], was lower than rates reported elsewhere in Cameroon and Myanmar [21, 46], in line with the 68.9% reported in Southwest Ethiopia [42] and higher compared to rates in Madagascar, Uganda and a host of eight other African countries [33, 35, 51].

Access to LLINs in the household of 69.1% in this study was lower compared to results reported elsewhere [21, 33, 42, 45], higher than the 21% reported in Batwa [43], within 57.3 – 78.8% in eight African countries [35] and 32.3 – 81.3% reported in a multi-country study [52]. The low household universal coverage and accessibility in this study could be attributed to the significant differences amongst the health districts and differences in family size vs gender of household head.

### Household utilisation of LLINs

Household universal LLINs utilisation of 20.5% (16.0% of the *de facto* population) was very low compared to previous studies elsewhere in and out of Cameroon [19, 20, 30, 33, 37, 45, 46, 48, 51, 53, 54]. This was however high compared to the 6.9 – 15.3% reported in Myanmar [32]. The very low household LLINs utilisation could be attributed to the significant differences amongst the health districts as well as household composition and the installation of LLINs on all beds in the household. It could also be due to inadequate education on LLINs utilisation, socio-political tensions, and differences in the different study designs.

Bednet utilisation by all children 0 – 5 years and expectant mothers in the household of 14.6% and 63.4%, respectively, is low compared to 63% vs 60% reported in BHD [19], 52% vs 58% in the national territory and elsewhere in the world [31, 33, 34, 37, 46, 51, 53, 55]. The low LLIN utilisation by all children 0 – 5 years old could be attributed to significant differences in the health districts, age and gender of household heads, educational status of the household head as well as the presence or absence of bushes or water pools around dwellings.

Use of LLINs by the household head last night of 28.0% was low compared to 58.3% reported in Rural and Semi-Urban communities in the South West Region of Cameroon [8] and 47.2 – 63.8% elsewhere [30, 42]. Meanwhile, the regular use of LLINs of 38.7% was low compared to 48% reported in China [30]. The low use of LLINs last night by household heads and regular use of LLINs could significantly be attributed to differences in the health districts and age of the household heads.

LLIN misuse of 2.3 – 22.7% was also similar to the 18.2% reported in Mezam Division [17] and 21% in Kenya and Ethiopia [56, 57]. The use of LLINs for other purposes, other than the prevention of mosquito bites could be attributed to: inadequate education on utilisation, normal social behaviour, lack of good playgrounds, as 2.8% of the households admitted that children used bednets as football goal post nets.

### Care and maintenance of LLINs

Out of the 1,251 household heads sampled, 1,089 (87.1%) said LLINs can be washed, while 627 (50.1%) affirmed the recommended LLIN washing frequency of 2 – 4 times a year. The recommended LLIN wash frequency reported in this study was similar to the 52.0% reported in Kenya [56] and lower than the 68 – 87.3% reported worldwide [58-60]. The optimal LLIN washing frequency could be attributed to the age and occupation of the household head as well as the health district.

## RECOMMENDATION

The populations of the three health districts should be properly educated by community health workers and stakeholders on the regular utilisation of LLINs and by all household occupants.

## CONCLUSIONS

Our findings highlighted low rates of household universal coverage, accessibility, and utilisation indicators as well as maintenance amidst high ownership of at least one LLIN per household and the free MDC. Quelling the on-going socio-political crisis and scaling up efforts can lead to increased coverage which may systematically contribute to household universal utilisation and thus reduce malaria morbidity. Our finding that health districts are strongly associated with LLINs ownership, utilisation and maintenance suggests that MDCs should be complemented by education and behaviour change communication, emphasizing that malaria is transmitted by mosquito bites and it can be prevented by sleeping under LLINs.

## STRENGTHS AND LIMITATIONS OF THE STUDY

### Strengths

The data used in this study was collected by trained surveyors. The health district offices were consulted for the mapping of the HAs, quarters and census list of households used in the last MDC and Expanded Programme on Immunisation campaigns. The quality of data collected was assured through the multistage sampling strategy to minimize bias and pretesting of questionnaires.

### Limitations

This was a cross-sectional study, representing the snapshot of the population within the study period and does not show cause and effect since the predictor and outcome variables were measured simultaneously. Data was collected through self-reporting and thus there is a possibility of recall bias where the respondent provides socially acceptable answers. In this study, however, respondents were required to only recall whether they and occupants of their households slept under a LLIN the previous night, as well as the source and number of LLINs in the household. Ownership, utilisation, and maintenance of LLINs in the three health districts in 2017 and 2018 could not be attributed solely to the 2015-2016 MDC, as other sources of LLINs: ANC for pregnant women and gift from a relation, and our study design could not capture the contribution of each intervention.

## What is already known on this topic

- The proportion, as well as factors associated with LLINs ownership and utilisation have been investigated in some health districts in Cameroon and Africa.
- The ownership of LLINs, leads to increased utilization in some health districts in Cameroon and Africa.

## What this study adds

- Information on the care and maintenance of LLINs.
- Information on ownership, care, and maintenance of LLINs shall enhance and improve on the prevention of malaria in the health districts concerned.

## Authors’ contributions

Conceptualization and Methodology: FNC, PNF, and PNF. Field data collection: FNC, PNF, BMC, SFM, YEN, and PKJ. Data curation/ Statistical analysis: FNC, PNF, and PKJ. Original draft preparation: FNC, PNF, SFM, BMC, PKJ, and PNF. Review and editing of draft: FNC, PNF, PKJ, PNF, and ASE. Administration and supervision: CNF, ANT, PNF, and ASE. All authors read and approved the final manuscript. FNC, PNF, and PKJ contributed equally to this work.

## Acknowledgements

We are thankful to the heads of households who participated in this survey, the community health workers, as well as field assistants who worked under challenging field conditions to ensure the success of the study.

The authors also acknowledge that there is a preprint of this manuscript on www.biorxiv.org, with doi: https://doi.org/10.1101/465005.

## Data Availability

The dataset used for analysis in this study, is provided within the study, and is provided here as (**S1 file: Dataset**). A preprint with DOI: https://doi.org/10.1101/465005. of this study is also available at www.biorxiv.org.

### ABBREVIATIONS

95% C.I: 95% Confidence Interval
BHD: Bamenda Health District
HA: Health area
LLINs: Long-lasting insecticidal nets
OR: Odds Ratio
*p*: Significance value
SD: Standard Deviation
SHD: Santa Health District
THD: Tiko Health District
χ^2^: Chi square

## Ethical Approval

Ethical clearance was obtained from the IRB-FHS of the University of Buea.

## Supplementary Materials

S1 file: Dataset. (*Supplementary Materials*)

