## Supplementary material for "Ownership, Coverage, Utilisation and Maintenance of Long-lasting insecticidal nets in Three Health Districts in Cameroon: A Cross-Sectional Study": S1 Questionnaire (from another version of this preprint: version 2, 2021-02-20)

### HOUSEHOLD QUESTIONNAIRE ON THE OWNERSHIP, UTILISATION AND MAINTENANCE OF MOSQUITO BED NETS IN BAMENDA, SANTA AND TIKO HEALTH DISTRICTS

*Please answer all the questions by completely ticking (√) in the box to the left of the answer or filling the blank spaces provided.*

Health Area:___________________________, Quarter: _____________________________

| | | | | | |
|---|---|---|---|---|---|
| | Identification information: | | | | |
| | Date of data collection: | Unique Identification Number: | | | |
| | Health District: | ☐ Bamenda | ☐ Santa | ☐ Tiko | |
| | **SECTION A: OWNERSHIP OF MOSQUITO BED NETS** | | | | |
| **1a.** | Do you have mosquito bed net(s) in your household? | | ☐ Yes | ☐ No | |
| **1b.** | If Yes, how many? (*Please write the number, say: 0, 1, 2, 3, …*) | | | | |
| **2.** | If you have mosquito bed nets (MBNs) in your household, how did you acquire it (them)? (*Please select and tick; √, all that applies*) | | | | |
| | ☐ The 1st Mass Distribution Campaign<br>☐ The 2nd Mass Distribution Campaign<br>☐ Ante-Natal Clinic (ANC) | ☐ I bought<br>☐ From a Relation (*Brother, Sister, Aunt, Uncle, Parents, Friend*)<br>☐ None of these | | | |
| **3a.** | Did you acquire mosquito bed nets during the second MDC (2015)? | | ☐ Yes | ☐ No | |
| **3b.** | If No, Why? (*Please select and tick; √, all that applies*) | | | | |
| | ☐ I was not censured<br>☐ I was told by distributors that there was a shortage of MBNs<br>☐ I did not go for collection | ☐ I was late for collection<br>☐ Others (*Specify*): ___________________________________ | | | |
| | **SECTION B: USAGE OF MOSQUITO BED NETS** | | | | |
| **4.** | Are the mosquito bed net(s) hung on all the beds in each room? | | ☐ Yes | ☐ No | |
| **5.** | How many people in the household slept under the mosquito net last night? (*Please write the number of persons, say 0, 1, 2, 3, … years old, in the box below the option*). | Children 0 – 5 | Children 6 – 17 | Expectant mother | Persons ≥18 |
| | **a)** Slept home | | | | |
| | **b)** Used MBNs | | | | |
| **6.** | Did you sleep under the mosquito bed net (MBN) last night? | | ☐ Yes | ☐ No | |
| **7.** | In the past week, for how many nights were the nets used? (*Please write the number of nights: 0, 1, 2, 3, 4, 5, 6, 7*): | | | | |
| **8.** | If mosquito bed nets were not used every night, why not? (*Tick; √, all that apply*) | | | | |
| | ☐ It gives heat<br>☐ They were dirty<br>☐ No Mosquitoes<br>☐ The Coil/ Spray/ Repellent was used | ☐ Forgot<br>☐ Hard to hang<br>☐ Others (*Specify*): ___________________ | | | |
| **9a.** | Can mosquito bed nets be washed? | | ☐ Yes | ☐ No | |
| **9b.** | If Yes, how many times do you wash mosquito bed nets in one year? (*Please write the number of times in one year: 0, 1, 2, 3, …*) | | | | |
| **10.** | If you have mosquito net(s), how do you wash it (them)? (*Tick; √, all that apply*). | | | | |
| | ☐ Never washed it<br>☐ I use Savon<br>☐ I use detergent (*Omo, Blu, Saba*) | ☐ I use "coco" soap<br>☐ Others (*Specify*): ___________________ | | | |
| **11.** | How often do you use the mosquito bed net(s)? (*Tick; √, only one*). | | | | |
| | ☐ I do not use it<br>☐ During the rainy season | ☐ Some of the nights<br>☐ Every night | | | |
| **12.** | How often do you tug the net(s) to the bed(s) when brought down in the evening? | | | | |
| | ☐ I don't use it<br>☐ Some of the nights | ☐ Every night (*Please select and tick; √, all that applies*) | | | |

| | | | | |
|---|---|---|---|---|
| **13.** | If you have mosquito net(s), from who did you learn how to use and maintain it (them)? (*Tick; √, all that apply*). | | | |
| | ☐ I have not learned yet<br>☐ From mass media (*Television*, *Radio*, *Newspaper*, *Internet*)<br>☐ From the agents of mass distribution campaign<br>☐ From hospital staff<br>☐ Others (*Neighbours*, *Friends*, *Relatives*) | | | |
| **14.** | How many of the available bed nets do you not use? (*Write the number*) | | | |
| **15a.** | Apart from using bed nets to protect from mosquito bites, are there any other uses around your quarter? | | ☐ Yes | ☐ No |
| **15b.** | If Yes, which other uses? (*Please select and tick; √, all that applies*). | | | |
| | ☐ Fishing<br>☐ Chicken shed<br>☐ Bathing shelter | ☐ Wall material<br>☐ Wire mesh on windows<br>☐ Others (*Specify*): ___________________ | | |
| **16.** | Do you know any other method for malaria prevention? | | ☐ Yes | ☐ No |
| **17.** | In your opinion, which is (are) the best way(s) to prevent malaria? (*Write in the spaces provided*). | | | |
| | A<br>B | | | |
| **18.** | In your household, what is the main method used to protect against malaria? (*Tick; √, only one*). | | | |
| | ☐ Keep compound clean<br>☐ Use antimalarial<br>☐ Use MBNs | ☐ Vaccination<br>☐ I don't know | | |
| | **SECTION C: SOCIO-DEMOGRAPHIC DATA** | | | |
| **19.** | Age (*Please write your age or date of birth*): | | | |
| **20.** | Gender (*Please tick; √, only the one that applies*): | ☐ Male | ☐ Female | |
| **21.** | What is your Marital Status? (*Please tick; √, only the one that applies*) | | | |
| | ☐ 1. Single<br>☐ 2. Widow(er)<br>☐ 3. Cohabitating | ☐ 4. Divorced<br>☐ 5. Separated<br>☐ 6 Married | | |
| **22.** | What was the **highest** level of schooling that you completed? (*Please tick; √, only the one that applies*). | | | |
| | ☐ 1. No formal education<br>☐ 2. Primary education<br>☐ 3. Secondary<br>☐ 4. Tertiary education (*After high school; HND, Bachelor, Master, PhD*). | | | |
| **23.** | What is your occupation? (*Please tick; √, only the one that applies*) | | | |
| | ☐ Civil Servant<br>☐ Private Sector<br>☐ CDC Worker<br>☐ House Wife<br>☐ Student | ☐ Business<br>☐ Farming<br>☐ Fishing<br>☐ Others (*Specify*): _______________________ | | |
| **24.** | House type (*Please tick; √, only the one that applies*) | | | |
| | ☐ Cement Block<br>☐ Mud block | ☐ Caraboat<br>☐ Make shift (Mixed). | | |
| **25.** | How many sleeping rooms are there in the house? (*Write the number*) ____________ | | | |
| **26.** | Which of these do you have around the household? (*Tick; √, all that apply*). | | | |
| | ☐ Bushes/ Farm/ Garden | ☐ Standing waters | ☐ None of these | |
| **27.** | How many times did people in the household suffer from malaria last year? (*Please tick; √, only the one that applies*) | | | |
| | ☐ 0. No time<br>☐ 1. One time a year<br>☐ 2. Two times a year<br>☐ 3. Three times a year | ☐ 4. Four times a year<br>☐ 5. Five times a year<br>☐ 6. Six times a year | | |

Thank you for your Patience and Cooperation

Please return the completed questionnaire to the Investigator
